# Cold Shock Fail to Restrain Pre-formed *Vibrio parahaemolyticus* Biofilm

**DOI:** 10.1101/529925

**Authors:** Wenying Yu, Qiao Han, Xueying Song, Jiaojiao Fu, Haiquan Liu, Zhuoran Guo, Pradeep K Malakar, Yingjie Pan, Yong Zhao

**Author notes:** These authors contributed equally to this study (Wenying Yu, Qiao Han).

## Abstract

The source of persistent infections can be biofilms that occur naturally on food surfaces and medical biomaterials. Biofilm formation on these materials are likely to be affected by environmental temperature fluctuations and information on noticeable temperature shifts on the fate of pre-formed biofilm is sparse. Changes to pre-formed *Vibrio parahaemolyticus* biofilm under cold shock (4 °C and 10 °C) was explored in this study. We show that *V. parahaemolyticus* biofilm biomass increased significantly during this cold shock period and there was a gradual increase of polysaccharides and proteins content in the extracellular polymeric matrix (EPS). In addition, we demonstrate that the expression of flagella and virulence-related genes were differentially regulated. The architecture of the biofilm, quantified using mean thickness (MT), average diffusion distance (ADD), porosity (P), biofilm roughness (BR) and homogeneity (H) also changed during the cold shock and these parameters were correlated (P < 0.01). However, the correlation between biofilm architecture and biofilm-related genes expression was relatively weak (P < 0.05). Cold shock at 4 °C and 10 °C is not sufficient to reduce *V. parahaemolyticus* biofilm formation and strategies to reduce risk of foodborne infections should take this information into account.

## Introduction

A biofilm is a multicellular complex, formed from microorganisms that are attached to a surface and generally embedded in extracellular polymeric substances (EPS) [1, 2]. Polysaccharides and proteins are the most prevalent components of EPS in biofilms as the production of mature biofilms requires polysaccharides to hold the cells together which maintain the structural integrity of the biofilm [3,4]. Biofilm is a means of persistence for bacteria and plays a crucial role in the bacterial life cycle [5]. Almost 65 % of all reported bacterial infections are caused by bacterial biofilms according to the Center for Disease Control and Prevention in the United States [6]. Bacterial cells embedded in biofilms are also more resistant towards disinfectants and antibiotics when compared to their free living or planktonic forms [7,8]. However, bacterial biofilm formation can be directly affected by environmental temperature [9,10] especially rapid decrease in temperature (cold shock).

*Vibrio parahaemolyticus* is a halophilic, gram-negative, curved rod-shaped bacterium [11] which is widely distributed in aquatic reservoirs. *V. parahaemolyticus* is frequently isolated from a variety of seafoods, particularly shellfish [12,13], and can persist on food or food-contact surface by forming biofilms [14]. These pathogenic bacteria are also leading causes of seafood-derived illness in many Asian countries, including China, Japan, and Korea [15–17] and disease outbreaks are highly correlated with temperature fluctuation [18,19].

Growth of planktonic *V. parahaemolyticus* is minimal or decreasing at temperatures below 10 °C [20,21]. However, Costerton et al. [22] reported that bacteria in biofilms are more adaptable to low temperatures than their free counterparts. Han et al. [23] found that *V. parahaemolyticus* at 4 °C and 10 °C formed monolayer biofilms and these biofilms were significantly different to biofilms formed at higher temperatures (15 °C - 37 °C). These low temperatures are typically applicable for transport, retail and processing of commercial seafoods [21,24] and information on noticeable temperature shifts on the fate of pre-formed *V. parahaemolyticus* biofilm is sparse.

Bacterial quorum sensing (QS) and the production of flagella, type III secretion systems (T3SS) and haemolysins (TDH and TRH) are closely coupled to biofilm formation of *V. parahaemolyticus*. Quorum sensing (QS) is a bacterial cell–cell communication process mediated by signaling molecules called autoinducers (AIs) [25] and essential for bacterial biofilm formation [26,27]. The autoinducers regulate the production of transcription factors (AphA and OpaR). AphA is an activator of virulence and biofilm formation in *V. parahaemolyticus* and OpaR represses biofilm formation in the pandemic O3:K6 *V. parahaemolyticus* [28]. The expression of *pilA* (chitin-regulated pilus pilin gene) contributes to flagella production, flagella are critical during early stages of bacterial colonization of a surface. In addition, the expression of genes involved in encoding the type III secretion systems (T3SS1 and T3SS2), the thermostable direct haemolysin (TDH) and the TDH-related haemolysin (TRH) correlates with increased biofilm production [29,30].

To date, little information is available on pre-formed bacterial biofilm changes at low-temperature shifts, although it is crucial for improving food safety and controlling bacterial infections outbreaks. *V. parahaemolyticus* is a widely distributed foodborne pathogen, temperature plays a great role in its survival. Researchers generally assume that cold environment can restrain biofilm formation and bacterial activity. This study explored the effects of *V. parahaemolyticus* biofilm upon a shift from 37 °C to 4 °C or 10 °C from two aspects. On the one hand, the changes of biofilm biomass and EPS contents, the expression of biofilm related genes directly described that pre-formed bacterial biofilm could not be controlled efficiently in cold environment. On the other hand, the CLSM images revealed biofilm morphological structure change, the correlation analysis showed inner relationship among biofilm structure parameters and biofilm related genes. According to previous investigations on cold tolerance of *V. parahaemolyticus*, one hypothesis was proposed that cold chain is essential to maintain food quality [31], but cold-chain transportation and low-temperature preservation are not such effective to control the bacteria in seafood when it has been exposed to the high temperature (15 - 37°C). For a certain time, allowing the establishment of a permanent bacterial community organized in biofilms. It is a potential risk for food safety. This study also provided useful information for ensuring seafood safety after refrigeration for food industry in the future.

## Materials and methods

### Bacterial strains and culture conditions

*V. parahaemolyticus* strain (ATCC17802) was inoculated from storage at − 80 °C into thiosulfate citrate bile salts sucrose agar (TCBS agar, Land Bridge Technology, Beijing, P.R.China) and incubated overnight at 37 °C. Single colony on the TCBS agar was cultured into 9 mL of Tryptic Soy Broth (TSB, Land Bridge Technology, Beijing, P.R.China) containing 3 % (w/v) NaCl and incubated overnight at 37 °C with shaking at 200 rpm. The *V. parahaemolyticus* cultures were diluted with fresh TSB (3 % NaCl) to an OD_600_ value of 0.4 for subsequent experiments.

### Biofilm formation assay

Static biofilms were grown in 24-well polystyrene microtiter plates (Sangon Biotech Co., Ltd., Shanghai, P.R.China) and biofilm production was measured using crystal violet staining with some modification [32]. In brief, the growth of biofilms was initiated by inoculating 10 μL of the cell suspension which were cultured in 2.1 into 990 μL of fresh TSB medium (3 % NaCl) in the individual wells. All samples were statically (without shaking) incubated at 37 °C for 24 h to obtain the pre-formed biofilm, then the pre-formed biofilm was shifted to low temperature (4 °C, 10 °C) or kept at 37 °C for 12 h, 24 h, 36 h, 48 h and 60 h without shaking, respectively. To prevent the medium evaporation, all plates were sealed with plastic bag. At each of these time points, the biofilms were quantified by crystal violet staining. Specifically, the suspension was discarded and the wells were gently washed three times with 1 mL of 0.1 M phosphate buffer (PBS) to remove non-adhered cells, and subsequently the biofilm was stained with 1 mL of 0.1% (w/v) crystal violet (Sangon Biotech Co., Ltd., Shanghai, P.R.China) for 30 min, then washed three times with 1 mL of 0.1 M PBS to remove unbound crystal violet. After drying for 45 min in a 60 °C incubator, biofilm stained by crystal violet was dissolved in 2 mL of 95% ethanol (Sinopharm Chemical Reagent Co., Ltd., Shanghai, P.R.China) for 30 min. Wells containing un-inoculated TSB served as negative controls, and the difference between the optical density of tested strains and negative control (OD_570_) was used to characterize the biofilm-forming ability of the tested strains [33]. This experiment was tested in triplicate.

### EPS analysis

The exopolysaccharides in *V. parahaemolyticus* biofilm were extracted using sonication method with some modification as described previously [34,35]. Biofilm cells on the wells were removed by vortexing and scraping after addition of 1 ml 0.01 M KCl (Sinopharm Chemical Reagent Co., Ltd., Shanghai, P.R.China). The cells were disrupted with a sonicator (JY92-IIN, Ningbo scientz biotechnology Co., Ltd., Ningbo, P.R.China) for 4 cycles of 5 s of operation and 5 s of pause. The sonicated suspension was centrifuged (4,000 *g*, 20 min, 4 °C), and the supernatant was filtered through a 0.22 μm membrane filter (Sangon Biotech Co., Ltd., Shanghai, P.R.China), 100 μL of the filtrate was mixed with 200 μL 99.9 % sulfuric acid in new tubes. After incubation at room temperature for 30 min, 6 % phenol was added to the mixture. Then after incubation at 90 °C for 5 min, the absorbance of mixture was measured at 490 nm. The amount of carbohydrate was quantified by dividing OD_490_ / OD_595_ values.

The amount of proteins was quantified by the Lowry method [35]. 40 μL of the filtrate was mixed with 200 μL Lowry reagent (L3540, Sigma Aldrich, St. Louis, Missouri, USA) in new tubes. After incubation at room temperature for 10 min, 20 μL Folin-Ciocalteu reagent (L3540, Sigma Aldrich, St. Louis, Missouri, USA) was added to the mixture. Then after incubation at room temperature for 30min, absorbance was measured at 750 nm. The amount of proteins was quantified by dividing OD at 750 nm by OD at 595 nm.

### Confocal laser scanning microscopy imaging

Quantitative parameters describing biofilm physical structure have been extracted from three-dimensional CLSM images and used to compare biofilm structures, monitor biofilm development, and quantify environmental factors affecting biofilm structure. *V. parahaemolyticus* biofilm was observed using a confocal laser scanning microscopy (CLSM). The biofilm was immobilized using 2 mL of 4 % (v/v) glutaraldehyde solution for 2 h at 4 °C, and rinsed 3 times with 0.1 M PBS and stained with SYBR Green I (Sangon Biotech Co., Ltd., Shanghai, P.R.China) for 30 min in darkness at room temperature [36]. The wells were then washed with 0.1 M PBS to remove the excess stain and air dried. CLSM images were acquired using a Zeiss LSM710-NLO Confocal Laser Scanning Microscopy (Carl Zeiss, Jena, Germany) with a 20× objective. Biofilm structure morphology in three dimensions was analyzed using Image Structure Analyzer-2 (ISA-2) [37,38].

### RNA extraction, reverse transcription, and RT-PCR analysis

Total RNA from biofilms were extracted and purified using the RNA extraction kit (Generay Biotech Co., Ltd., Shanghai, P.R.China), according to the manufacturer's instructions. RNA concentrations were determined by measuring the absorbance at 260 nm and 280 nm (ND-1000 spectrophotometer, NanoDrop Technologies, Wilmington, DE, USA). Reverse transcription (RT) was performed with 200 ng total RNA using the Prime Script RT reagent Kit with gDNA Eraser (Takara, Dalian, P.R.China) following the manufacturer’s instructions.

The qPCR reaction mixture (20 μl) contained 10 μl mix, 0.4 μL ROXⅡ, 0.8 μM of the appropriate forward and reverse PCR primers, 2 μl of template cDNA and ddH_2_O 6μL. The reactions were preformed using an Applied Biosystems 7500 Fast Real-Time PCR System (Applied Biosystems, Carlsbad, USA). Negative controls (deionized water) were included in each run. Amplifications were performed in duplicate. The expression levels of all of the tested genes were normalized using the 16S rRNA gene as an internal standard [39]. Relative quantification was measured using the 2^−ΔΔCt^ method (the amount of target, normalized to an endogenous control and relative to a calibrator, where ΔΔCt = (Ct target − Ct reference) sample − (Ct target − Ct reference) calibrator) [40]. The specific primers (Table 1) were designed by Primer Premier 5.0 software, and all synthesized by Shanghai Sangon Biotech Company.

**Table 1.**
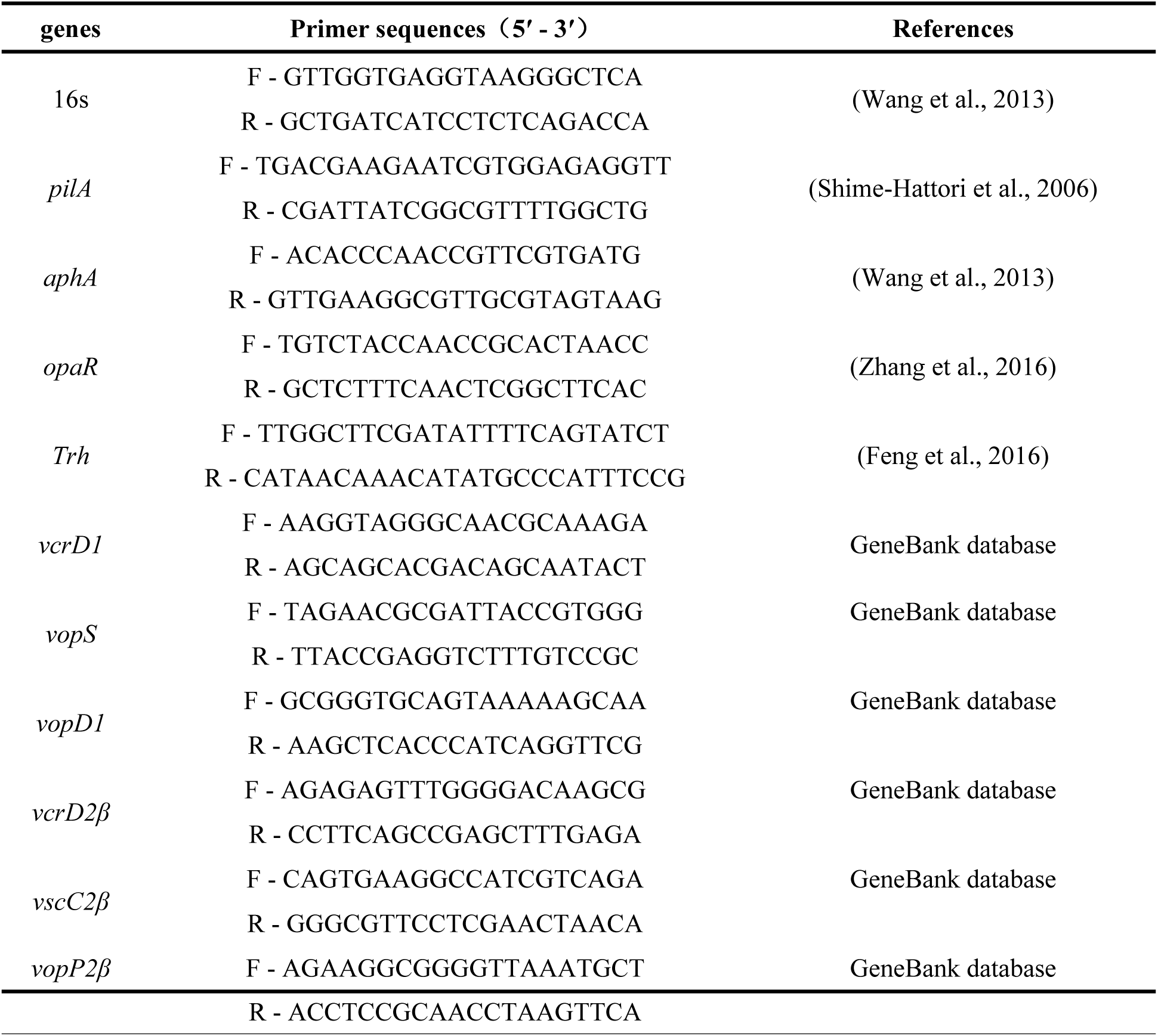
Primer sequences of the RT-qPCR assay.

### Statistical analysis

The statistical analysis was performed using the SPSS statistical software (version 17.0; SPSS Inc., Chicago, IL, USA), including two-way Analysis of Variance (ANOVA) for time-course evaluations, the Student *t*-test for comparison between groups, Pearson correlation coefficient at the 0.01 and 0.05 significant level. Values were considered significantly different if *p* < 0.05. Calculations and figures were performed using Microsoft Excel 2007 (Microsoft Corporation, Redmond, WA, USA) and origin 8.0, respectively. Linear regression analysis using Excel 2007.

## Results

### Biofilm biomass changes

The biomass changes of biofilm obtained using crystal violet staining under cold shock conditions is illustrated in Fig 1. Overall, regardless of the exposure to the cold shock, biofilm biomass increased with incubation time (Fig 1). At 4 °C, 10 °C and 37 °C, the starting OD_570nm_ of biofilm was 0.399 (pre-formed at 37 °C for 24h) and the OD_570nm_ continually increased of which 37 °C is significantly higher than 4 °C and 10 °C.

**Fig 1.**
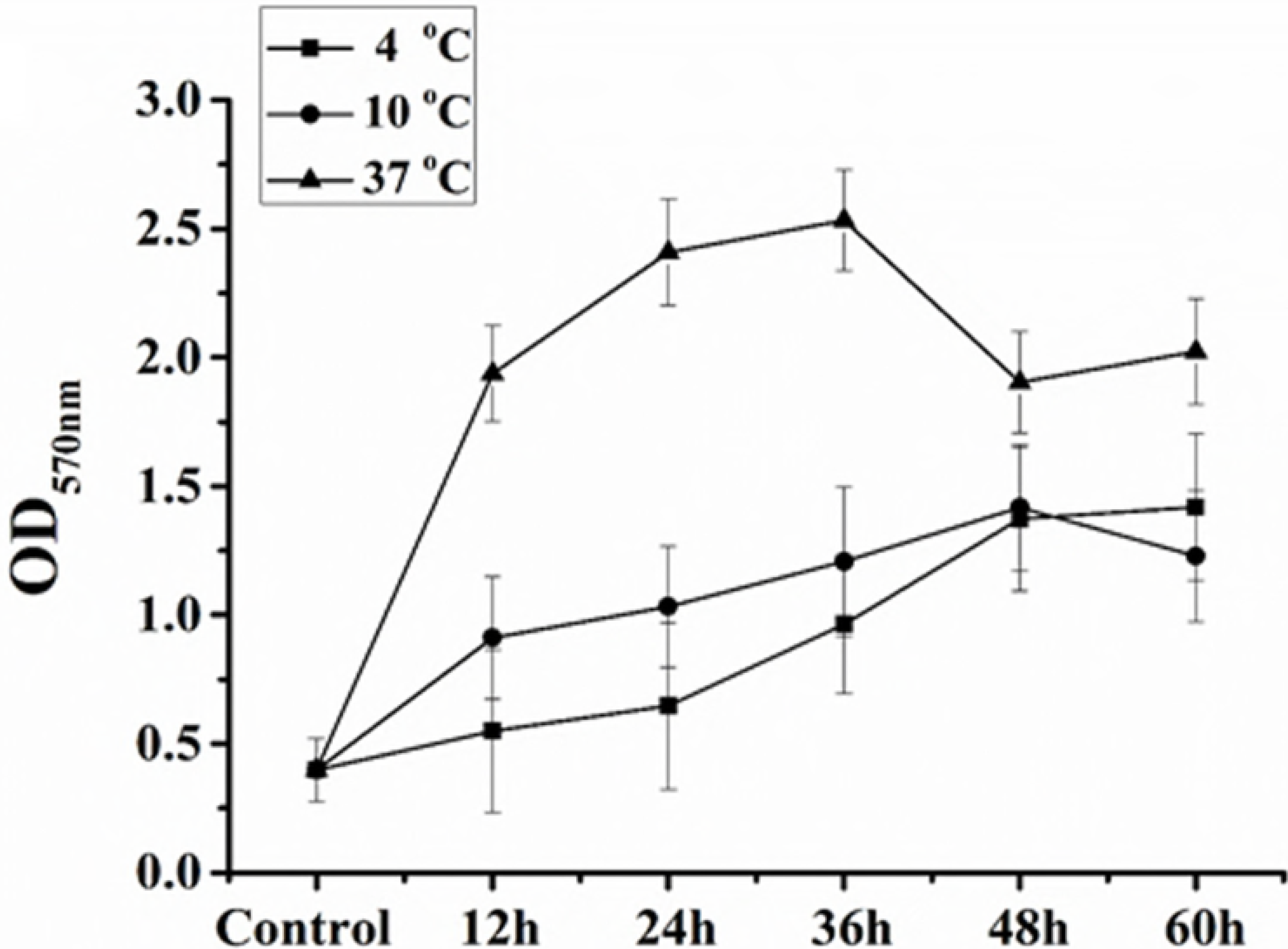
The changes of biofilm biomass. The change of biofilm biomass of *V. parahaemolyticus* exposed to cold shock (4 °C and 10 °C) or kept at 37 °C. The data are presented as the mean of OD _570nm_ ± standard deviation for three independent replicates.

### EPS analysis

To evaluate the effects of cold shock on EPS production, we analyzed the total exopolysaccharides and proteins in EPS of the pre-formed biofilm treated from 12 h to 60 h. As shown in Fig 2, the exopolysaccharides contents increased, and no remarkable difference of exopolysaccharides contents between 4 °C and 10 °C. We also observed that a higher (p < 0.05) exopolysaccharides production at 37 °C for 24 h and 36 h culture in comparison with that at low temperature shock. However, when treated for 48h, exopolysaccharides contents in EPS exposed to cold shock were higher than that at 37 °C. The results demonstrated that cold shock could only reduce the biosynthesis of exopolysaccharides, rather than restrain it absolutely, which coincident with the results from the crystal violet staining. Fig 2C shows the correlation between protein and polysaccharides in biofilm. The linear regressions of the data are.

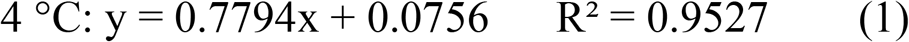

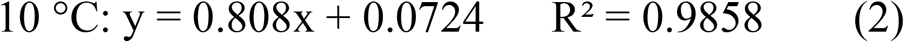

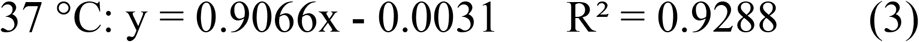

The higher R^2^ indicates a higher degree of linear dependence and the consistency of protein and polysaccharides. Meanwhile, The Pearson correlation coefficients were 0.976, 0.993 and 0.964 at 4 °C, 10 °C and 37 °C, respectively. The correlations of protein and polysaccharides at the three temperature conditions were all significant at the 0.01 level (2-tailed). Those results indicated that the compositions of EPS are high consistency. With the increase of biofilm biomass, the compositions of EPS are more coordinate and the order of biofilm structure is stronger than 37 °C.

**Fig 2.**
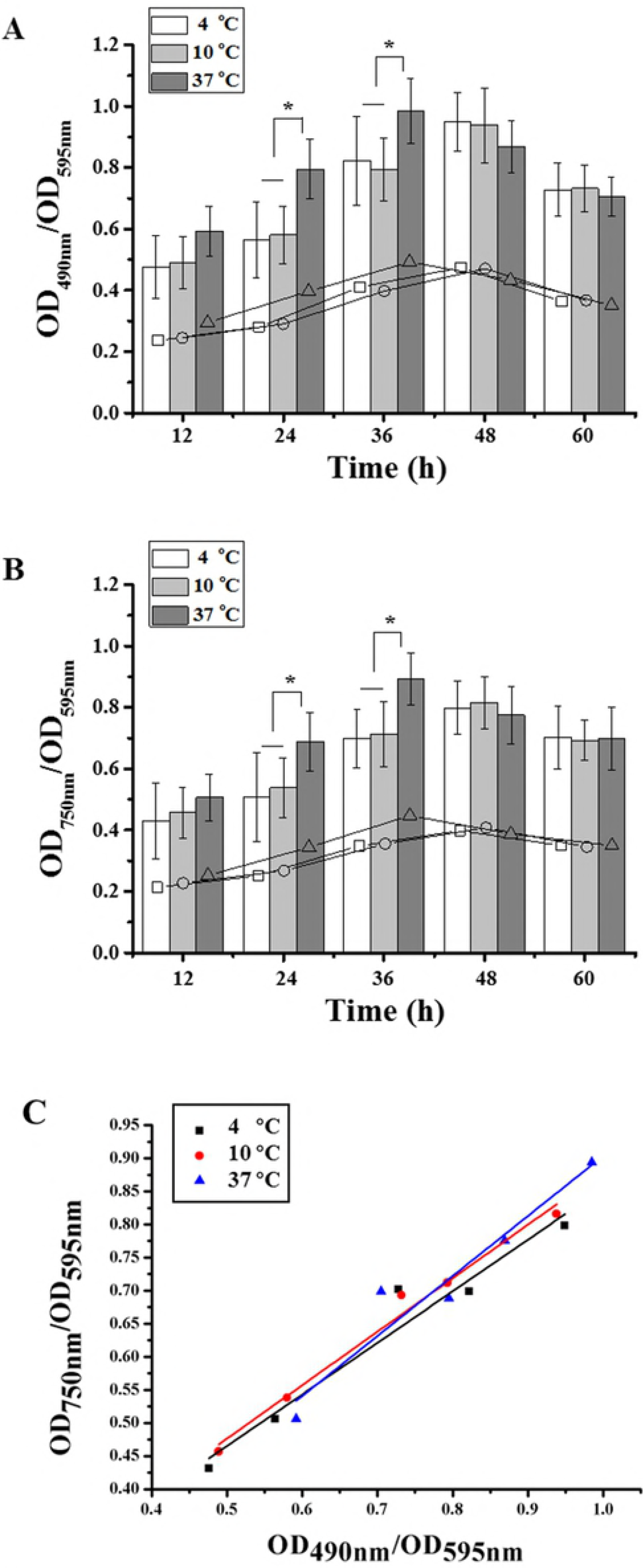
The contents change of EPS constituent. The contents change of EPS constituent exposed to cold shock (4 °C and 10 °C) or kept at 37 °C. (A) Total contents (column) of exopolysaccharides in EPS of *V. parahaemolyticus* biofilm, and the average value (line) which showed the variation tendency of exopolysaccharides after cold shock. (B) Total contents (column) of proteins in EPS of *V. parahaemolyticus* biofilm, and the average value (line) which showed the variation tendency of proteins after cold shock. Error bars represent the standard deviations of 3 measurements. * indicates significant difference (p < 0.05). (C) The linear regression of total contents of exopolysaccharides and proteins in EPS of *V. parahaemolyticus* biofilm.

### Gene expression analysis

In order to better understand the fate of pre-formed biofilm of *V. parahaemolyticus* exposed to cold shock, we further analyzed the biofilm-related gene expression changes, including genes encoding for flagella (*pilA*), QS *(aphA*, *opaR*), virulence (*trh*) and T3SS (*vcrD1*, *vopS*, *vopD1*, *vscC2β*, *vcrD2β*, *vopP2β*). As shown in Fig 3, all of the selected flagella and virulence genes were differentially expressed in the biofilm cells. Also, with the increase of incubation time, the genes *aphA* and *vscC2β* and *vopP2β* were up regulated gradually, whereas the genes *opaR* and *vopS* were expressed without significant difference. However, after cold shock, T3SS genes (*vcrD1*, *vcrD2β* and *vopD1*) were down-regulated. Earlier studies showed that *vcrD1* and *vcrD2β* encodes for an inner membrane protein [41], and *vopD1* is essential for translocation of T3SS1 effector of *V. parahaemolyticus* [42]. Compared with 4 °C and 10 °C, the genes expression of flagella (pilA), QS (aphA, opaR) and virulence (trh) at 37 °C are significantly higher. Additionally, genes involved in T3SS genes (particularly *vcrD1*, *vopD1* and *vcrD2β*) downregulated more obvious at 37 °C.

The results showed that, although *V. parahaemolyticus* biofilm grow better at constant 37 °C, the biofilm cells have adapted to the low temperature shift. This is consistent with the findings of biofilm biomass changes and EPS changes.

**Fig 3.**
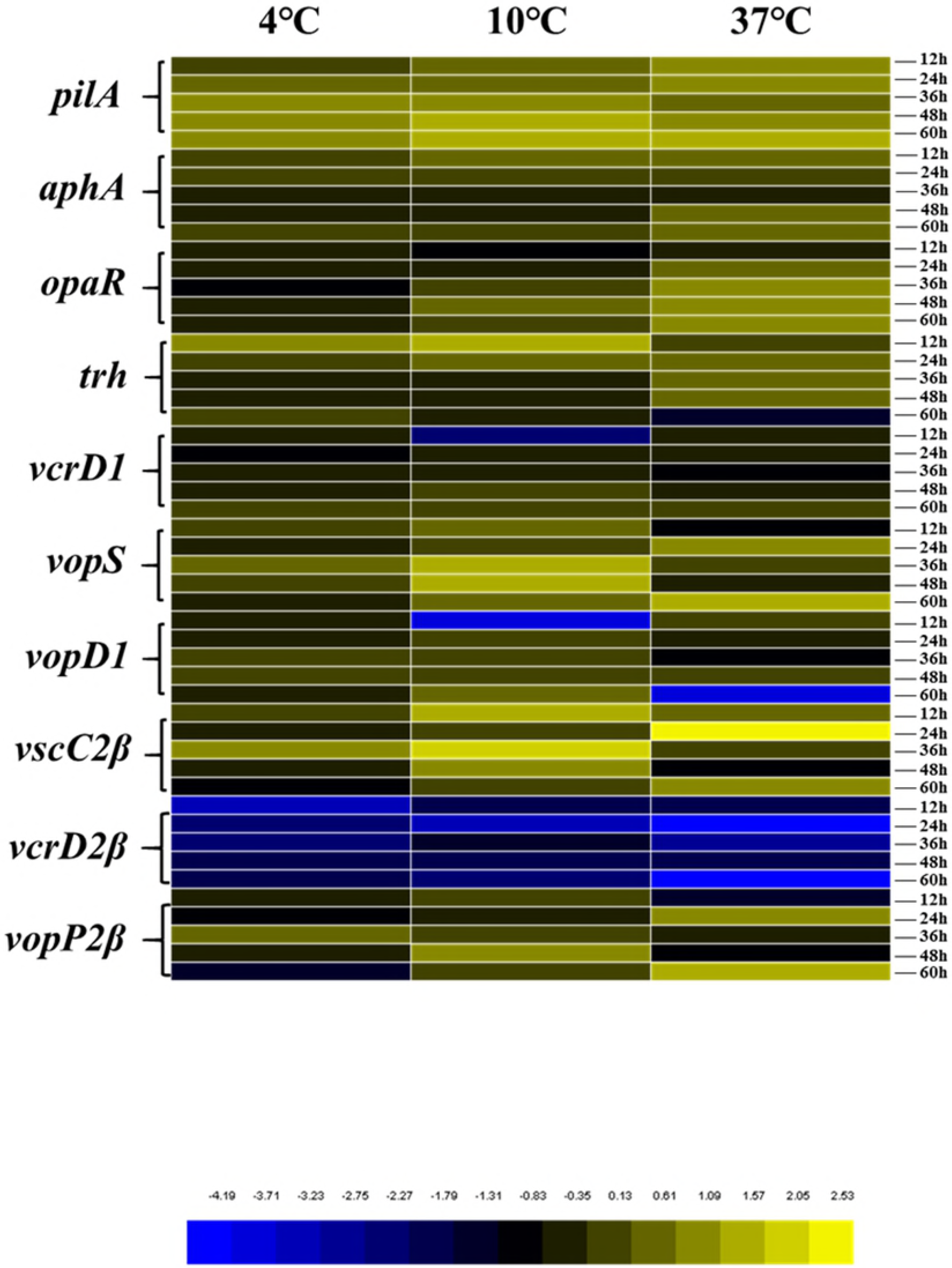
Expression profiles of *V. parahaemolyticus* biofilm related genes. Expression profiles of a selected set of genes in *V. parahaemolyticus* biofilm exposed to 4 °C and 10 °C or kept at 37 °C for 12h, 24h, 36h, 48h and 60h. Induced expression is represented in yellow, repressed expression is represented in blue, and little changed expression is represented in black. Differential expression of genes involved in flagella, QS, virulence gene, T3SS1 and T3SS2 were observed upon a cold shift. The color scale is shown at the lower right corner. Primers sequences used in RT-qPCR assay are provided in Table1.

### Biofilm architecture changes

The CLSM images of pre-formed *V. parahaemolyticus* biofilm are presented in Fig 4 A and shows that *V. parahaemolyticus* is able to form complex three-dimensional structures. Compared with pre-formed biofilm (Fig 4A), the biofilm thickness at three conditions (4 °C, 10 °C and 37 °C) have actually increased. Fig 4B shows that with increasing incubation time, the biofilm architecture changed from a flat homogeneous layer of cells to a complex structure. Specially, the contact surfaces were completely covered by dense and homogeneous biofilm when cultured at 4 °C and 10 °C for 36 h or at 37 °C for 24 h.

**Fig 4.**
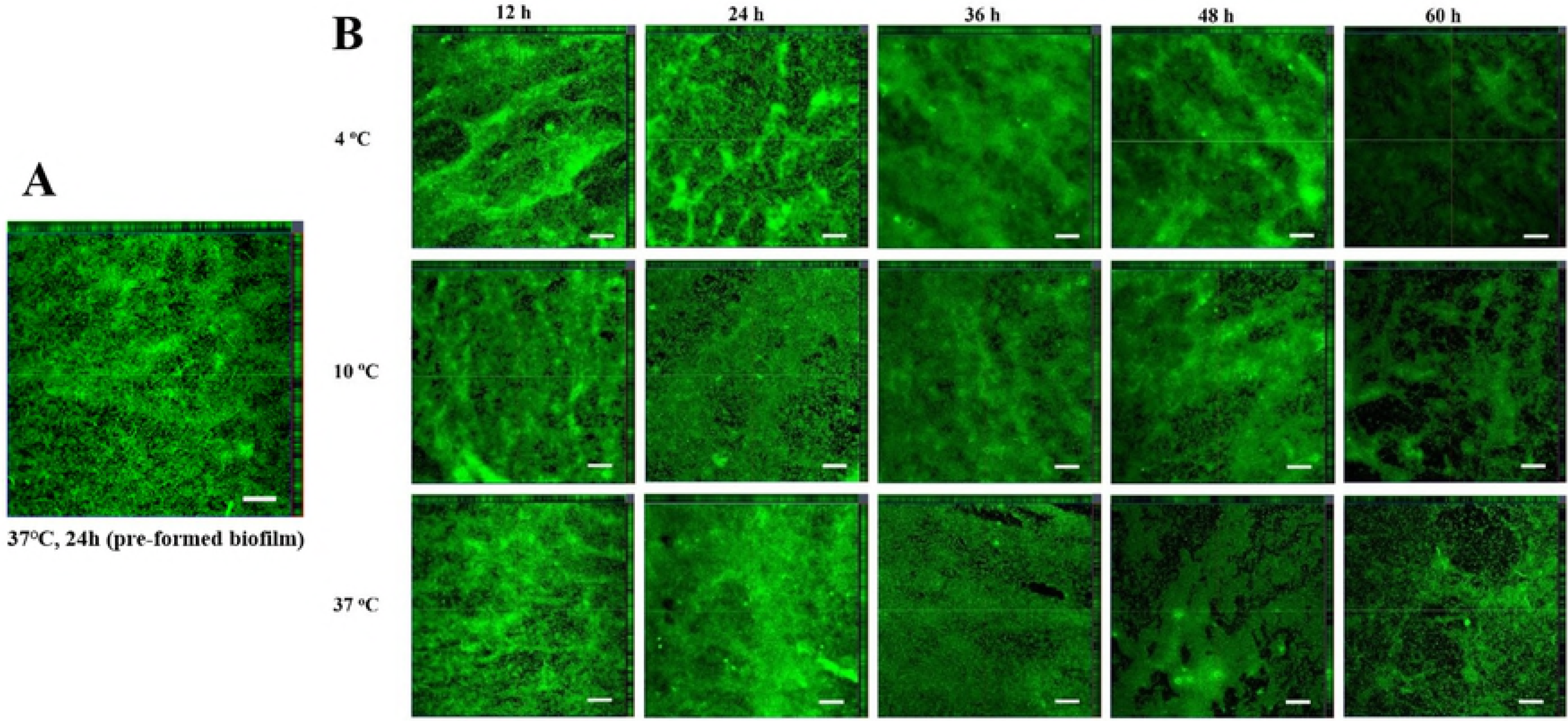
Confocal laser scanning microscopy (CLSM) images. (A) Confocal laser scanning microscopy (CLSM) images of pre-formed biofilm formed by *V. parahaemolyticus* at 37 °C for 24 h. The scale bar represents 20 μm. (B) Confocal laser scanning microscopy (CLSM) images of biofilm formed by *V. parahaemolyticus* subjected to cold shock (4 °C and 10 °C) or kept at 37 °C. The biofilms were incubated at 37 °C for 24 h to obtain preformed biofilm, and then shifted to 4 °C and 10 °C immediately or kept at 37 °C for 12 h, 24 h, 36 h, 48 h and 60 h. The scale bar represents 20 μm.

Quantitative parameters describing biofilm physical structure are summarized in Fig 5. The parameter MT provides a measure of the spatial size of the biofilm and is the most common variable used in biofilm literature. The MT value of pre-formed biofilm was 1.77 and changed slightly when exposed to cold shock at 10 °C, while the changes at 4 °C and 37 °C were significant. Average diffusion distances (ADD) have been suggested as a measurement of the distance over which nutrients and other substrate components diffused from the voids to the bacteria within micro-colonies [43,44]. The initial ADD value of pre-formed biofilm in this study was 1.04 and the ADD of biofilm at the low temperatures (4 and 10 °C) were higher than that at 37 °C.

**Fig 5.**
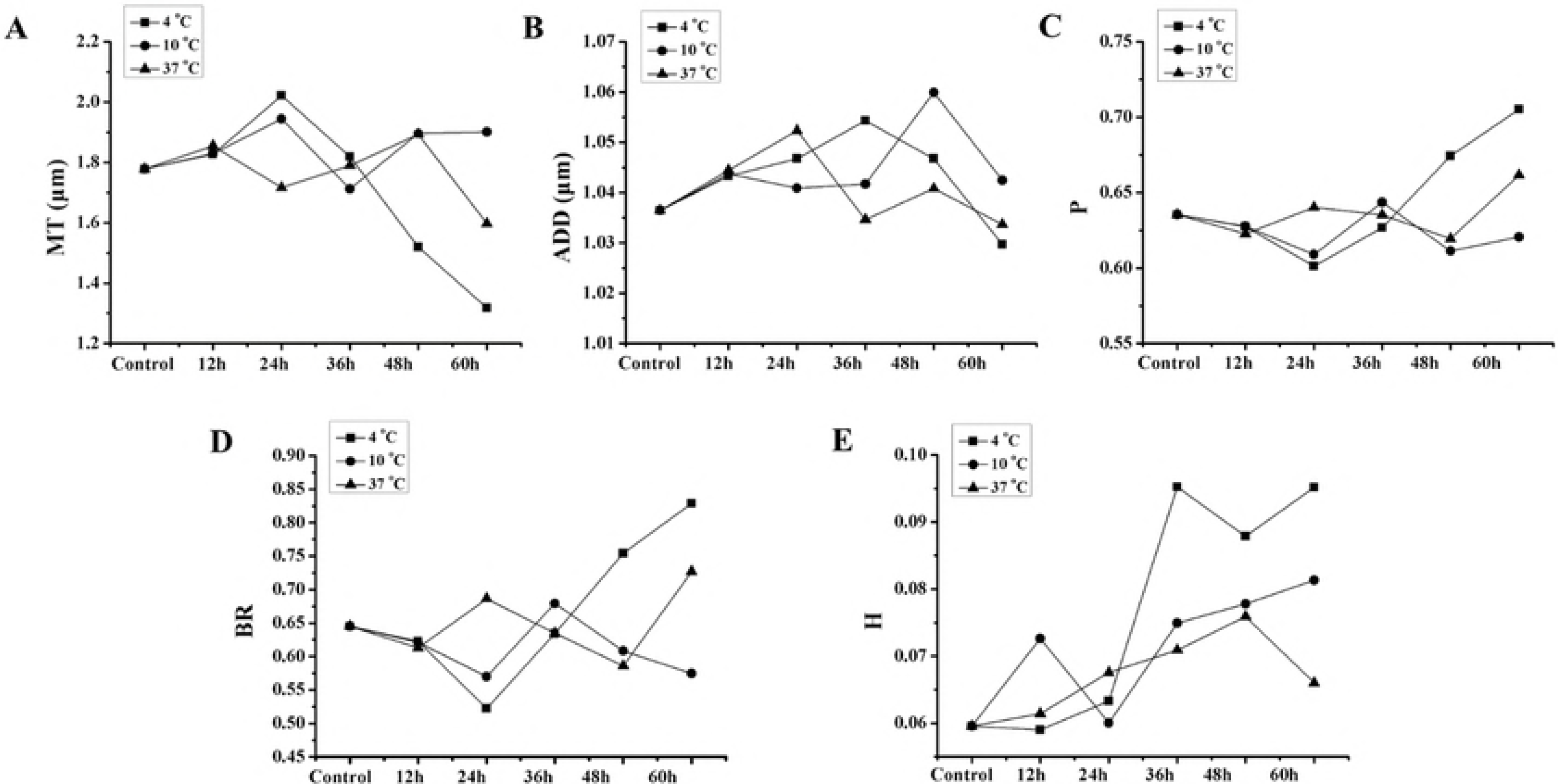
The changes of biofilm structural parameters. Quantification of structural parameters changes (3A-E) in biofilm formed by *V. parahaemolyticus* after low temperature shift, including mean thickness (MT), average diffusion distance (ADD), porosity (P), biofilm roughness (BR) and homogeneity (H).

Biofilm roughness (BR) is a good measure for the variability in biofilm thickness [45] and the BR value of pre-formed biofilm in this study was 0.64. The BR value significantly increased after cold shock (4 °C), (Fig 5D) while the BR value at 37 °C showed no significant change during the incubation time. Areal parameters describe the morphological structures of biofilm and we selected porosity (P) as a measure of this parameter where the porosity decreases with increasing number of cell clusters. The initial porosity value of pre-formed biofilm was 0.64 and Fig 5C shows that after cold shock (4 °C), the porosity value was significantly increased, in the range from 0.601 to 0.705, with the increasing of incubation time. Textural entropy is a measure of randomness in the gray scale of the image and the higher the textural entropy, the more heterogeneous the image is. When shifted to low temperatures, the pre-formed biofilm needed to accommodate the changed circumstances to form mature biofilm. The CLSM images were in accordance with the results of crystal violet staining. These results indicated that cold shock could only prolonged the period of biofilm mature.

### Correlation analysis

The correlations among biofilm structure parameters are listed in Table 2. Analysis of the 6 biofilm structure parameters of *V. parahaemolyticus* at 4, 10 and 37 °C showed a positive correlation between biofilm thickness (BR) and porosity (P). The spatial sizes of the biofilm (MT) and biofilm thickness (BR) were negatively correlated as well as spatial size of the biofilm (MT) and porosity (P). Compared with 10 °C, those trends are more obvious at 4 °C.

**Table 2.** The correlation among biofilm structural parameters.

| Temperature<br>(°C) | Biofilm<br>structural<br>parameters | MT |  | BR |  | P |  | ADD |  | H |  |
| --- | --- | --- | --- | --- | --- | --- | --- | --- | --- | --- | --- |
|  |  | A | B | A | B | A | B | A | B | A | B |
| 4 | MT | 0 | 1 | **0.000** | **-0.996\*\*** | **0.000** | **-0.999\*\*** | 0.196 | 0.692 | 0.204 | -0.683 |
|  | BR |  |  | 0 | 1 | **0.001** | **0.992\*\*** | 0.243 | -0.642 | 0.182 | 0.707 |
|  | P |  |  |  |  | 0 | 1 | 0.177 | -0.713 | 0.222 | 0.664 |
|  | ADD |  |  |  |  |  |  | 0 | 1 | 0.863 | -0.108 |
|  | H |  |  |  |  |  |  |  |  | 0 | 1 |
| 10 | MT | 0 | 1 | **0.007** | **-0.968\*\*** | **0.008** | **-0.966\*\*** | 0.729 | 0.215 | 0.640 | -0.287 |
|  | BR |  |  | 0 | 1 | 0.055 | 0.870 | 0.987 | -0.01 | 0.697 | 0.240 |
|  | P |  |  |  |  | 0 | 1 | 0.497 | -0.406 | 0.600 | 0.320 |
|  | ADD |  |  |  |  |  |  | 0 | 1 | 0.540 | 0.370 |
|  | H |  |  |  |  |  |  |  |  | 0 | 1 |
| 37 | MT | 0 | 1 | **0.001** | **-0.992\*\*** | 0.601 | -0.319 | 0.719 | 0.222 | 0.590 | 0.328 |
|  | BR |  |  | 0 | 1 | 0.723 | 0.220 | 0.875 | -0.098 | 0.525 | -0.382 |
|  | P |  |  |  |  | 0 | 1 | 0.197 | -0.691 | 0.736 | 0.209 |
|  | ADD |  |  |  |  |  |  | 0 | 1 | 0.745 | -0.201 |
|  | H |  |  |  |  |  |  |  |  | 0 | 1 |

Note: A, Sig. (2-tailed); B, Correlation coefficient; \*, correlation is significant at the 0.05 level (2-tailed); \*\*, correlation is significant at the 0.01 level (2-tailed).

Table 3 displays the correlation between biofilm formation related genes and biofilm structural parameters at 4, 10 and 37 °C. There were appreciable correlations between *pilA* and H; *vopP2β* and MT, BR at the 0.05 level (2-tailed); *vcrD1* and MT, BR, P; *vscC2β* and ADD. In conclusion, after cold shock, the flagella and T3SS genes have significant correlation with biofilm structure in *V. parahaemolyticus*.

**Table 3.**
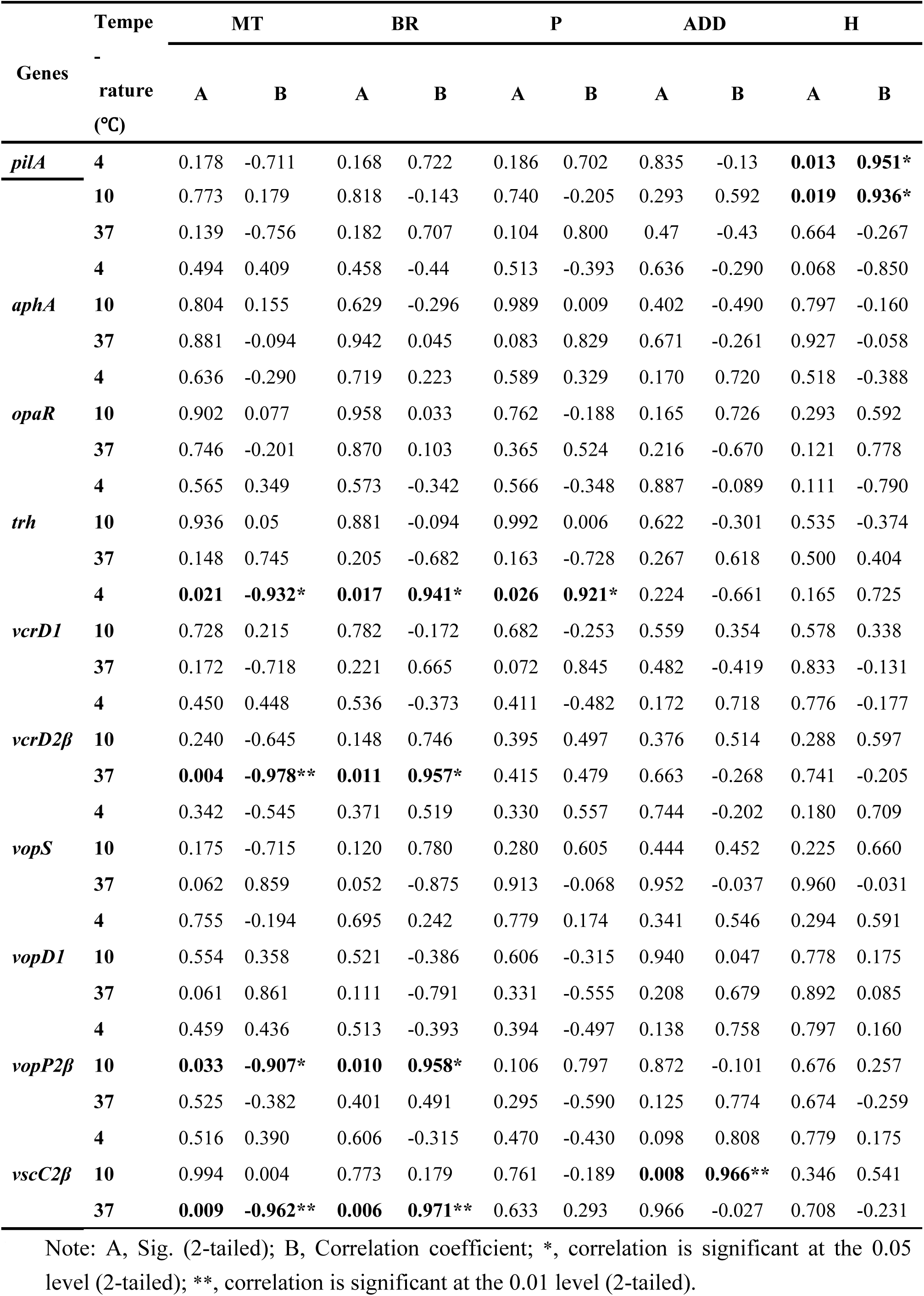
The correlation between biofilm structural parameters and biofilm formation related genes.

## Discussion

Pathogenic *V. parahaemolyticus* poses a serious threat to public health, and the majority of cases are associated with the consumption of seafood contaminated with this pathogen. Since biofilms shields the encased cells from chemical sanitizers and environmental stress, the biofilm-forming capability of *V. parahaemolyticus* contributes to its persistence and transmission in the environment [46]. Previous studies concentrated on biofilm formation at constant temperatures [10,47], where in reality the food environment fluctuates.

In this study, we gained further insights into the fate of pre-formed *V*. *parahaemolyticus* biofilm under cold chain temperatures. The *V*. *parahaemolyticus* cells in the biofilm gradually adapted to cold shock and protected themselves against the cold shift. Fig 1 showed that biofilm biomasses were increased when shifted to low temperature (4 °C and 10 °C). The genes encoding for the production of flagella, virulence and QS (*aphA*) were up-regulated after cold shock, thus improving the chance of survival of the biofilm cells. This up-regulation of the molecular machinery also contributes to biofilm structure. As it shown in Table 3, the *V. parahaemolyticus* flagella and T3SS genes (*pilA*, *vcrD1*, *vopP2β* and *vscC2β*) have significant correlation with the biofilm structural parameters under cold shift. Flagella in *Vibrio spp*. is intricately involved in attachment to surfaces, usually by enhancing movement towards the surface [4, 48]. Enos-Berlage et al. [49] examined biofilm formation of *V*. *parahaemolyticus* and observed that *flgE* and *flgD* mutants were defective in attachment and biofilm formation. Asadishad et al. [50] also found that bacterial swimming motility was decreased and the transcription of flagellin encoding genes were expressed in *Bacillus subtilis* exposed to cold temperature. We speculate that a similar mechanism operates during *V. parahaemolyticus* biofilm formation.

The bacterial T3SS nanomachine is a virulence mechanism which delivers effector proteins directly from the bacterial cytosol to host cells [51,52]. However, several studies concluded that the expression of T3SS genes were repressed in biofilm-growing bacteria [53]. Ferreira et al. [54] demonstrated that T3SS1 genes expression and biofilm production were inversely regulated. In this study, we found that genes encoding T3SS (*vcrD1*, *vopD1* and *vcrD2β*) were also significantly down regulated after cold shock. The results were consistent with the research of Ferreira et al. and Kuchma et al. [54,55] which indicated that the expression of T3SS genes can inversely regulate biofilm formation of *V. parahaemolyticus*. Yet despite the connection, we interestingly found T3SS (*vopS*, *vscC2β* and *vopP2β*) were up-regulated. This suggests that the relationship between biofilm formation and T3SS genes expression varied by the exact T3SS genes.

In T3SS mutants, the adherence to surfaces during biofilm formation was impaired and the expression of proteins involved in metabolic processes, energy generation, EPS production and bacterial motility as well as outer membrane proteins were also impacted [55]. The change of EPS production after cold shock may be linked to the regulation of T3SS genes and Jennings et al. [56] reported that the T3SS encoded by *Salmonella* Pathogenicity Island 1 (SPI1) mediates biofilm-like cell aggregation. Therefore, the flagellum encoding gene and the T3SS gene are known to be involved in regulating biofilm formation at low temperature and these mechanisms helps *V. parahaemolyticus* adapt the adverse environment.

The cold chain is a primary factor in the preservation and transportation of perishable foods and ensures that consumers can enjoy safe and good quality foods. However, previous studies showed that the efficiency of the cold chain was often less than ideal [57]. In this study, we demonstrated that cold shock induced the expression of genes involved in biofilm formation and increased biofilm biomass continuously, causing that the biofilm cells had the ability to adapt to the low temperature shift. It is interesting to investigate how biofilm cells to sense the cold shock signal and regulate the gene expression in further research. From the grow trends of *V. parahaemolyticus* biofilm biomass, polysaccharide and protein content, the expression of biofilm-related genes, it is possible that the biofilm begins to dissolute after 60h for the limitation of environmental resource which also need a further research. We recommend additional research on the molecular mechanisms between biofilm formation and low temperature fluctuations, especially the role of the membrane proteins in sensing environmental temperature, such as cold-shock proteins (CSPs) and cold acclimation proteins (Caps). These proteins may be overexpressed during prolong growth of the cold-tolerant bacteria.

## Conclusions

In summary, we demonstrate that cold shock (4 °C and 10 °C) induces changes of gene expression related with biofilm formation and biofilm structure, reliance on cold shock for reducing risk of foodborne infections should take this information into account.

## Acknowledgments

This research was supported by the National Natural Science Foundation of China (31671779 and 31571917), National Key R&D Program of China (2018YFC1602200 and 2018YFC1602205)， Shanghai Agriculture Applied Technology Development Program (Grant No. G20160101 and T20170404), Innovation Program of Shanghai Municipal Education Commission (2017-01-07-00-10-E00056), the “Dawn” Program of Shanghai Education Commission (15SG48).

## References

1. Hall-Stoodley L, Costerton JW, Stoodley P. Bacterial biofilms: from the natural environment to infectious diseases. Nature Reviews Microbiology. 2004; 2 (2): 95–108. https://doi.org/10.1038/nrmicro821. PMID: 15040259

2. Janissen R, Murillo DM, Niza B, Sahoo PK, Nobrega MM, Cesar CL, et al. Spatiotemporal distribution of different extracellular polymeric substances and filamentation mediate Xylella fastidiosa adhesion and biofilm formation. Scientific Reports. 2015; 5: 9856. https://doi.org/10.1038/srep09856. PMID: 25891045

3. Li T, Bai R, Liu J. Distribution and composition of extracellular polymeric substances in membrane-aerated biofilm. Journal of Biotechnology. 2008; 135 (1): 52–57. https://doi.org/10.1016/j.jbiotec.2008.02.011. PMID: 18403037

4. Yildiz FH, Visick KL. Vibrio biofilms: so much the same yet so different. Trends in Microbiology. 2009; 17 (3): 109–118. https://doi.org/10.1016/j.tim.2008.12.004. PMID: 19231189

5. Sultana M, Nusrin S, Hasan NA, Sadique A, Ahmed KU, Islam A, et al. Biofilms Comprise a Component of the Annual Cycle of Vibrio cholerae in the Bay of Bengal Estuary. Mbio. 2018; 9(2): e00483–18. https://doi.org/10.1128/mBio.00483-18. PMID: 29666284

6. Potera C. Microbiology - Forging a link between biofilms and disease. Science. 1999; 283 (5409): 1837–1839. https://doi.org/10.1126/science.283.5409.1837. PMID: 10206887

7. Caraher E, Reynolds G, Murphy P, McClean S, Callaghan M. Comparison of antibiotic susceptibility of Burkholderia cepacia complex organisms when grown planktonically or as biofilm in vitro. European Journal of Clinical Microbiology & Infectious Diseases. 2007; 26 (3): 213–216. PMID: 17265071

8. Elexson N, Afsah-Hejri L, Rukayadi Y, Soopna P, Lee HY, Tuan Zainazor TC, et al. Effect of detergents as antibacterial agents on biofilm of antibiotics-resistant Vibrio parahaemolyticus isolates. Food Control. 2014; 35 (1): 378–385. https://doi.org/10.1016/j.foodcont.2013.07.020.

9. Di Ciccio P, Vergara A, Festino AR, Paludi D, Zanardi E, Ghidini S, et al. Biofilm formation by staphylococcus aureus, on food contact surfaces: relationship with temperature and cell surface hydrophobicity. Food Control. 2015; 50: 930–936. https://doi.org/10.1016/j.foodcont.2014.10.048.

10. Pan YW, Breidt FJ, Gorski L. Synergistic effects of sodium chloride, glucose, and temperature on biofilm formation by listeria monocytogenes serotype 1/2a and 4b strains. Applied & Environmental Microbiology. 2010; 76(5): 1433. https://doi.org/10.1128/AEM.02185-09.. PMID: 20048067

11. Wong HC, Ting SH, Shieh WR. Incidence of toxigenic vibrios in foods available in Taiwan. The Journal of applied bacteriology. 1992; 73: 197–202. https://doi.org/10.1111/j.1365-2672.1992.tb02978.x. PMID: 1399913

12. Yeung PS, Boor KJ. Epidemiology, Pathogenesis, and Prevention of Foodborne Vibrio parahaemolyticus Infections. Foodborne Pathog Dis Foodborne Pathogens and Disease. 2004; 1: 74–88. PMID: 15992266

13. Vora GJ, Meador CE, Bird MM, Bopp CA, Andreadis JD, Stenger DA. Microarray-based detection of genetic heterogeneity, antimicrobial resistance, and the viable but nonculturable state in human pathogenic Vibrio spp. Proceedings of the National Academy of Sciences of the United States of America. 2005; 102 (52): 19109–19114. https://doi.org/10.1073/pnas.0505033102. PMID: 16354840

14. Annous BA, Smith JL, Fratamico PM, Solomon E. B. 20 – Biofilms in fresh fruit and vegetables. Biofilms in the Food and Beverage Industries. 2009; 517–535.https://doi.org/10.1533/9781845697167.4.517.

15. Su YC, Liu C. Vibrio parahaemolyticus: a concern of seafood safety. Food Microbiology. 2007; 24 (6): 549–558. https://doi.org/10.1016/j.fm.2007.01.005. PMID: 17418305

16. Chao GX, Jiao XN, Zhou XH, Wang F, Yang ZQ, Huang JL, et al. Distribution of genes encoding four pathogenicity islands (VPaIs), T6SS, biofilm, and type I pilus in food and clinical strains of Vibrio parahaemolyticus in China. Foodborne Pathogens and Disease. 2010; 7 (6): 649. https://doi.org/10.1089/fpd.2009.0441. PMID: 20132020

17. Broberg CA, Calder TJ, Orth K. Vibrio parahaemolyticus cell biology and pathogenicity determinants. Microbes and Infection. 2011; 13 (12-13): 992–1001. https://doi.org/10.1016/j.micinf.2011.06.013. PMID: 21782964

18. Duan J, Su YC. Comparison of a Chromogenic Medium with Thiosulfate‐Citrate‐Bile Salts‐Sucrose Agar for Detecting Vibrio parahaemolyticus. Journal of Food Science. 2005; 70 (2): M125–M128. https://doi.org/10.1111/j.1365-2621.2005.tb07102.x.

19. Turner JW, Malayil L, Guadagnoli D, Cole D, Lipp EK. Detection of Vibrio parahaemolyticus, Vibrio vulnificus and Vibrio cholerae with respect to seasonal fluctuations in temperature and plankton abundance. Environmental Microbiology. 2013; 16 (4): 1019–1028. https://doi.org/10.1111/1462-2920.12246. PMID: 24024909

20. Cook DW and Ruple AD. Indicator bacteria and Vibrionaceae multiplication in post-harvest shellstock oysters. Journal of Food Protection. 1989; 52: 343–349. https://doi.org/10.4315/0362-028X-52.5.343.

21. Burnham VE, Janes ME, Jakus LA, Supan J, DePaola A, Bell J. Growth and survival differences of Vibrio vulnificus and Vibrio parahaemolyticus strains during cold storage. Journal of Food Science. 2009; 74 (6): M314–318 PMID: 19723217.

22. Costerton JW, Stewart PS, Greenberg EP. Bacterial Biofilms: A Common Cause of Persistent Infections. Science. 1999; 284 (5418): 1318–1322. https://doi.org/10.1126/science.284.5418.1318. PMID: 10334980

23. Han N, Mizan MFR, Jahid IK, Ha SD. Biofilm formation by Vibrio parahaemolyticus on food and food contact surfaces increases with rise in temperature. Food Control. 2016; 70: 161–166. https://doi.org/10.1016/j.foodcont.2016.05.054.

24. US Food and Drug Administration (FDA). Quantitative risk assessment on the public health impact of pathogenic Vibrio parahaemolyticus in raw oysters. Center for Food Safety and Applied Nutrition, Food and Drug Administration, US Department of Health and Human Services. 2005.

25. Ng WL, Bassler BL. Bacterial quorum-sensing network architectures. Annu Rev Genet Annual Review of Genetics. 2009; 43: 197–222. https://doi.org/10.1146/annurev-genet-102108-134304.

26. Watnick PI, Fullner KJ, Kolter R. A role for the mannose-sensitive hemagglutinin in biofilm formation by Vibrio cholerae El Tor. Journal of Bacteriology. 1999; 181 (11): 3606 PMID: 10348878.

27. Sun FJ, Zhang YQ, Wang L, Yan XJ, Tan YF, Guo ZB, et al. Molecular Characterization of Direct Target Genes and cis-Acting Consensus Recognized by Quorum-Sensing Regulator AphA in Vibrio parahaemolyticus. Plos One. 2012; 7 (9): e44210. https://doi.org/10.1371/journal.pone.0044210. PMID: 22984476

28. Wang L, Ling Y, Jiang HW, Qiu YF, Qiu JF, Chen HP, et al. AphA is required for biofilm formation, motility, and virulence in pandemic Vibrio parahaemolyticus. International Journal of Food Microbiology. 2013; 160 (3): 245–251. https://doi.org/10.1016/j.ijfoodmicro.2012.11.004. PMID: 23290231

29. Paranjpye RN, Myers MS, Yount EC, Thompson JL. Zebrafish as a model for Vibrio parahaemolyticus virulence. Microbiology. 2013; 159: 2605–2615. https://doi.org/10.1099/mic.0.067637-0. PMID: 24056807

30. Calder T, de Souza Santos M, Attah V, Klimko J, Fernandez J, Salomon D, et al. Structural and regulatory mutations in Vibrio parahaemolyticus type III secretion systems display variable effects on virulence. FEMS Microbiology Letters. 2014; 361 (2): 107–114. https://doi.org/10.1111/1574-6968.12619. PMID: 25288215

31. Laguerre O, Hoang HM, Flick D. Experimental investigation and modelling in the food cold chain: thermal and quality evolution. Trends in Food Science & Technology. 2013; 29(2), 87–97.

32. Djordjevic D, Wiedmann M, McLandsborough LA. Microtiter Plate Assay for Assessment of Listeria monocytogenes Biofilm Formation. Applied and Environmental Microbiology. 2002; 68 (6): 2950–2958. https://doi.org/10.1128/AEM.68.6.2950-2958.2002. PMID: 12039754

33. Han Q, Song X, Zhang Z, Fu J, Wang X, Malakar PK, et al. Removal of Foodborne Pathogen Biofilms by Acidic Electrolyzed Water. Frontiers in Microbiology. 2017; 8: 988. https://doi.org/10.3389/fmicb.2017.00988. PMID: 28638370

34. Gong AS, Bolster CH, Benavides M, Walker SL. Extraction and analysis of extracellular polymeric substances: comparison of methods and extracellular polymeric substance levels in Salmonella pullorum SA 1685. Environmental Engineering Science. 2009; 26 (10): 1523–1532. https://doi.org/10.1089/ees.2008.0398.

35. Kim HS, Park HD. Ginger extract inhibits biofilm formation by Pseudomonas aeruginosa PA14. Plos One. 2013; 8 (9): e76106. https://doi.org/10.1371/journal.pone.0076106. PMID: 24086697

36. Song X, Ma Y, Fu J, Zhao A, Guo Z, Malakar PK, et al. Effect of temperature on pathogenic and non-pathogenic Vibrio parahaemolyticus biofilm formation. Food Control. 2017; 73: 485–491. https://doi.org/10.1016/j.foodcont.2016.08.041.

37. Beyenal H, Donovan C, Lewandowski Z, Harkin G. Three-dimensional biofilm structure quantification. Journal of Microbiological Methods. 2004; 59: 395–413. https://doi.org/10.1016/j.mimet.2004.08.003. PMID: 15488282

38. Resat H, Renslow RS, Beyenal H. Reconstruction of biofilm images: combining local and global structural parameters. Biofouling. 2014; 30 (9): 1141–1154. https://doi.org/10.1080/08927014.2014.969721. PMID: 25377487

39. Eleaume H, Jabbouri S. Comparison of two standardisation methods in real-time quantitative RT-PCR to follow Staphylococcus aureus genes expression during in vitro growth. Journal of Microbiological Methods. 2004; 59 (3): 363–370. https://doi.org/10.1016/j.mimet.2004.07.015. PMID: 15488279

40. Livak KJ, Schmittgen TD. Analysis of relative gene expression data using real-time quantitative PCR and the 2(-Delta Delta C(T)) Method. Methods. 2001; 25 (4): 402–408. https://doi.org/10.1006/meth.2001.1262. PMID: 11846609

41. Park KS, Ono T, Rokuda M, Jang MH, Okada K, Iida T, et al. Functional characterization of two type III secretion systems of Vibrio parahaemolyticus. Infection and Immunity. 2004; 72 (11): 6659–6665. https://doi.org/10.1128/IAI.72.11.6659-6665.2004. PMID: 15501799

42. Shimohata T, Mawatari K, Iba H, Hamano M, Negoro S, Asada S, et al. VopB1 and VopD1 are essential for translocation of type III secretion system 1 effectors of Vibrio parahaemolyticus. Canadian Journal of Microbiology. 2012; 58 (8): 1002–1007. https://doi.org/10.1139/W2012-081. PMID: 22827847

43. Lewandowski Z, Webb D, Hamilton M, Harkin G. Quantifying biofilm structure. Water Science and Technology. 1999; 39: 71–76. https://doi.org/10.1016/S0273-1223(99)00152-3. PMID: 29649702

44. Yang X, Beyenal H, Harkin G, Lewandowski Z. Quantifying biofilm structure using image analysis. Journal of Microbiological Methods. 2000; 39 (2): 109–119. https://doi.org/10.1016/S0167-7012(99)00097-4. PMID: 10576700

45. Heydorn A, Nielsen AT, Hentzer M, Sternberg C, Givskov M, Ersbøll BK, et al. Quantification of biofilm structures by the novel computer program COMSTAT. Microbiology. 2000; 146 (Pt 10): 2395. https://doi.org/10.1099/00221287-146-10-2395. PMID: 11021916

46. Newton AE, Garrett N, Stroika SG, Halpin JL, Turnsek M, Mody RK. Increase in Vibrio parahaemolyticus infections associated with consumption of Atlantic Coast shellfish--2013. Morbidity and Mortality Weekly Report. 2014; 63 (15): 335. PMID: 24739344

47. Kadam SR, den Besten HM, van der Veen S, Zwietering MH, Moezelaar R, Abee T. Diversity assessment of Listeria monocytogenes biofilm formation: impact of growth condition, serotype and strain origin. International Journal of Food Microbiology. 2013; 165 (3): 259–264. https://doi.org/10.1016/j.ijfoodmicro.2013.05.025. PMID: 23800738

48. O’Toole GA, Kolter R. Flagellar and twitching motility are necessary for Pseudomonas aeruginosa biofilm development. Molecular Microbiology. 1998; 30 (2): 295. https://doi.org/10.1046/j.1365-2958.1998.01062.x. PMID: 9791175

49. Enos-Berlage JL, Guvener ZT, Keenan CE, McCarter LL. Genetic determinants of biofilm development of opaque and translucent Vibrio parahaemolyticus. Molecular Microbiology. 2005; 55 (4): 1160–1182. https://doi.org/10.1111/j.1365-2958.2004.04453.x. PMID: 15686562

50. Asadishad B, Olsson AL, Dusane DH, Ghoshal S, Tufenkji N. Transport, motility, biofilm forming potential and survival of Bacillus subtilis exposed to cold temperature and freeze-thaw. Water Research. 2014; 58: 239–247. https://doi.org/10.1016/j.watres.2014.03.048. PMID: 24768703

51. Cornelis GR. The type III secretion injectisome. Nature Reviews Microbiology. 2006; 4:811–825. https://doi.org/10.1038/nrmicro1526. PMID: 17041629

52. Kudryashev M, Stenta M, Schmelz S, Amstutz M, Wiesand U, Castano-Diez D, et al. In situ structural analysis of the Yersinia enterocolitica injectisome. Elife. 2013; 2: e00792. https://doi.org/10.7554/eLife.00792. PMID: 23908767

53. Kuchma SL, Connolly JP, O’Toole GA. A three-component regulatory system regulates biofilm maturation and type III secretion in Pseudomonas aeruginosa. Journal of Bacteriology. 2005; 187 (4): 1441–1454. https://doi.org/10.1128/JB.187.4.1441-1454.2005. PMID: 15687209

54. Ferreira RB, Chodur DM, Antunes LC, Trimble MJ, McCarter LL. Output targets and transcriptional regulation by a cyclic dimeric GMP-responsive circuit in the Vibrio parahaemolyticus Scr network. Journal of Bacteriology. 2012; 194 (5): 914–924. https://doi.org/10.1128/JB.05807-11. PMID: 22194449

55. Zimaro T, Thomas L, Marondedze C, Sgro GG, Garofalo CG, Ficarra FA, et al. The type III protein secretion system contributes to Xanthomonas citri subsp. citri biofilm formation. BMC Microbiology. 2014; 14: 96. https://doi.org/10.1186/1471-2180-14-96. PMID: 24742141

56. Jennings ME, Quick LN, Ubol N, Shrom S, Dollahon N, Wilson JW. Characterization of Salmonella type III secretion hyper-activity which results in biofilm-like cell aggregation. Plos One 2012; 7 (3): e33080. https://doi.org/10.1371/journal.pone.0033080. PMID: 22412985

57. Mercier S, Villeneuve S, Mondor M, Uysal I. Time–temperature management along the food cold chain: a review of recent developments. Comprehensive Reviews in Food Science and Food Safety. 2017; 16(4).

